# Cryo-EM Structure of the Pol Polyprotein Provides Insights into HIV Maturation

**DOI:** 10.1101/2021.10.03.462959

**Authors:** Jerry Joe E. K. Harrison, Dario Oliveira Passos, Jessica F. Bruhn, Joseph D. Bauman, Lynda Tuberty, Jeffrey J. DeStefano, Francesc Xavier Ruiz, Dmitry Lyumkis, Eddy Arnold

## Abstract

Many retroviral proteins are initially translated from unspliced full-length RNA as polyprotein precursors that are subsequently processed by the viral protease (PR) to yield the mature forms. In HIV-1, the enzymes, PR, reverse transcriptase (RT), and integrase (IN), are produced as part of the Gag-Pol polyprotein. While structures of the mature proteins have aided our understanding of catalytic mechanisms and the design of antiretroviral drugs, knowledge of the architecture and functional implications of the immature forms prior to PR-mediated cleavage is limited. We developed a system to produce and purify the HIV-1 Pol polyprotein intermediate precursor and determined its high-resolution cryo-EM structure. The RT portion of the polyprotein has an architecture similar to the mature RT p66/p51 heterodimer, and dimerization of the RT portion draws together two PR monomers to activate proteolytic processing. HIV-1 thus may leverage the dimerization interfaces in Pol to regulate the assembly and maturation of the polyprotein precursors.

## Main

Many of the gene products of RNA viruses are initially translated as precursor polyproteins that are subsequently cleaved to yield mature proteins that play either structural or enzymatic roles^1^. Early association events can affect the functions of the polyproteins during viral assembly and maturation. The spatial and temporal regulation of assembly and processing can modulate enzymatic functions, association with other proteins, nucleic acids, cofactors, and substrates, as well as the subcellular localizations of the components during infection and virus particle formation^2,3^. In the case of HIV-1 and other retroviruses, premature processing can lead to incomplete assembly and maturation of virions. Given that polyproteins can be long-lived and have distinct sequences at their processing junctions, they may form distinct inhibitor-binding sites from the mature proteins for drug targeting, as exemplified by bevirimat blockage of CA– SP1 processing within Gag^4,5^.

In HIV-1, the structural proteins are translated initially as part of Gag polyprotein, while the enzymes that enable the virus to multiply and spread are synthesized as part of the Gag-Pol precursor polyproteins. Both polyproteins are cleaved by the viral protease (PR), which is synthesized as part of Gag-Pol, during maturation^6^. Gag is processed to yield the structural proteins of the mature virion, matrix (MA), capsid (CA), and nucleocapsid (NC). Pol is processed to produce the viral enzymes PR, reverse transcriptase (RT), and integrase (IN). Gag-Pol is synthesized by translational readthrough; different retroviruses employ different read-through mechanisms. HIV-1 uses a translational frameshift mechanism. The efficiency of the frameshifting creates a Gag:Gag-Pol ratio of 20:1^1,4,5,7^ making Gag-Pol a relatively minor constituent of the virion (∼100 copies per virion).

Proteolytic maturation is initiated by activation of the viral PR while it is still embedded in the Gag-Pol polyprotein. PR is an obligate homodimer and activation requires dimerization of at least the PR portion of two Gag-Pol molecules. In virions, the mechanism of PR activation and polyprotein processing has been challenging to analyze in large part because purified virions mature asynchronously^8^. In the absence of *in virio* data, studies with *in vitro* translated polyprotein suggest that activation of PR, which appears to be a slow step in virion maturation, requires cleavage events that occur at the C-terminal end of Gag and within the trans-frame region (TFR or p6*). The TFR/p6* is an unstructured linker of variable length (56-68 amino acid residues depending on type or clade) that connects Gag to Pol in the Gag-Pol polyprotein^9-13^. These studies show that the PR embedded in the liberated Pol continues processing polyprotein precursors with similar sluggishness until the mature PR is released, suggesting an architectural similarity of the PR that is embedded in Pol to that in Gag-Pol^14^. It is worth noting that HIV-1 Pol expressed separately, or as a PR-RT fusion [similar to the mature form of the protein found in prototype foamy virus (PFV)] retains PR and RT enzymatic activities both *in vitro* and *in virio*^15,16^. Thus, Pol provides the simplest model system with which to understand the early events of PR activation and virion maturation. A more thorough understanding of Pol and its architectural configuration would yield valuable insights into a poorly characterized aspect of the retroviral replication cycle.

We have a detailed understanding of the atomic-level organization of mature processed forms for both the HIV-1 structural proteins (MA, CA, NC) and the enzymes (PR, RT, IN), including wild-type and drug-resistant variants, and we have a good understanding of how they interact with other proteins, nucleic acid substrates, and inhibitors. These structures have helped in the development of antiretroviral drugs^17-19^. There is also useful information about the structures of the Gag polyprotein and the immature virion^20-25^. However, much less is known about the molecular organization of the Pol portion of the Gag-Pol polyprotein, or the immature forms of the viral enzymes. This is due, at least in part, to the absence of systems that can be used to efficiently produce and purify large amounts of the HIV-1 Gag-Pol or Pol polyproteins for structural and other biophysical studies. To address these limitations, we have developed a system for efficient production and purification of HIV Pol polyprotein and have determined the structure of the dimeric HIV-1 Pol. The architectural organization of the constituent enzymes helps to explain prior data on Pol activity and provide important insights into viral maturation.

## Results

### Expression and purification of the Pol Polyprotein

Proteolytic degradation of heterologously expressed proteins in bacteria is a common problem that has been difficult to mitigate with genetic approaches and/or media engineering^26^. We have discovered that proteolytic degradation of HIV-1 Pol (and other proteins) expressed in *E. coli* cells was highly attenuated in the presence of 50 mM or higher concentrations of Mg^2+^ and synergistically complemented by low pH (6.0 and below) using a modified recipe of the original lysogeny broth (LB)^27^ (see Methods, **Extended Data Fig. 1** and **Extended Data Table 1** for details). Degradation of the HIV-1 Pol polyprotein construct was monitored by Western blot analysis with the anti-IN Mab 8E5 antibody (**Extended Data Fig. 1**) while varying the composition of the growth media. An HIV-1 Pol construct from the BH10 strain that was found to have favorable properties for biochemical and structural analyses was further optimized through the incorporation of multiple mutations. First, we incorporated an inactivating D25A mutation in the PR, which was necessary to avoid auto-catalytic cleavage of the polyprotein. Second, amino acids L659/F660 were mutated to D/D at the RT/IN cleavage junction to reduce proteolytic degradation and improve solubility. Finally, we incorporated an Sso7d tag and an HRV14 3C protease cleavage site at the N-terminus of the p6* region (designated HIV-1 Pol-D25A and subsequently referred to as HIV-1 Pol) (**Fig. 1a**). HIV-1 Pol was purified using nickel affinity followed by HiTrap heparin chromatography, and finally gel filtration using a Superose 6 Increase 10/300 GL column (see Methods for details). The gel filtration elution peak contained a shoulder, suggesting the presence of structural heterogeneity (which could be either conformational or compositional) in the polyprotein (**Fig. 1b**). SDS-PAGE analysis revealed a single band (**Fig. 1c**), and dynamic light scattering (DLS) of the pooled fractions showed that 99.9% of the sample by mass was accounted for by a single species with a hydrodynamic radius of 6 nm, consistent with a single-chain multimeric protein (**Fig. 1d**). Thus, our optimized bacterial expression conditions and the multi-step purification procedure yielded suitable amounts (> 3 mg/L) of pure protein amenable for structural and biochemical studies.

**Figure 1.**
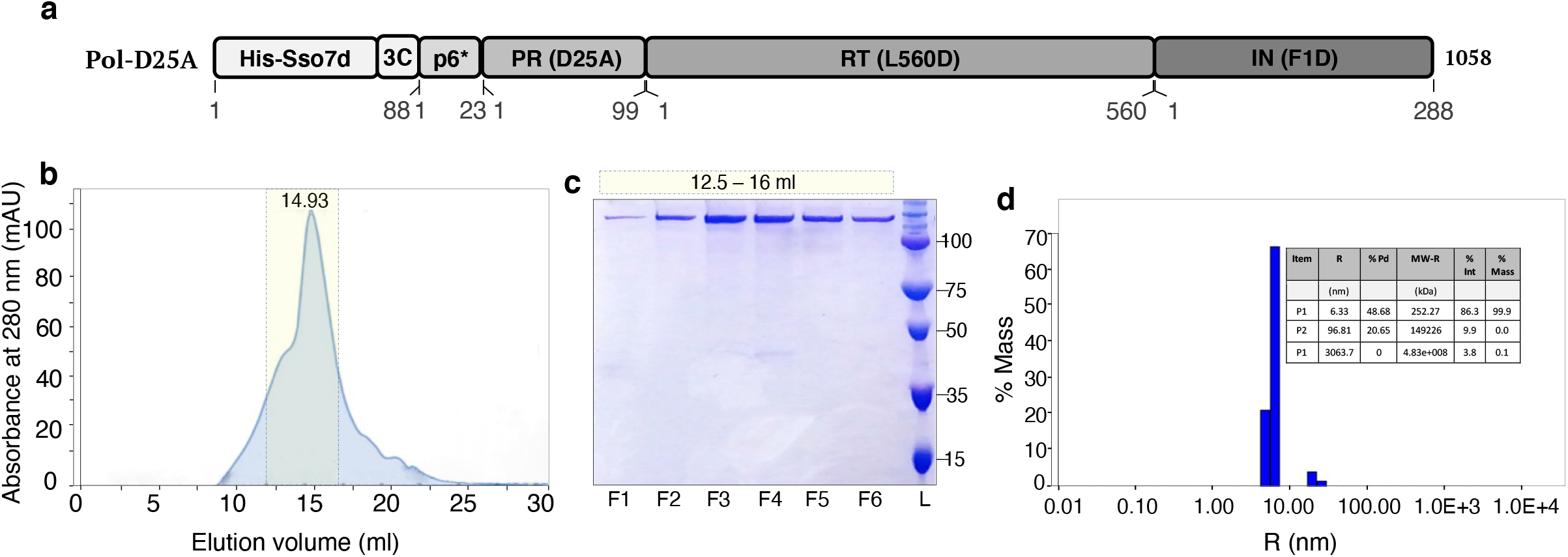
Expression and biophysical characterization of HIV-1 Pol. **a**, Annotated schematic of the HIV-1 Pol construct used for this study. **b**, Superose 6 Increase gel filtration profile of purified HIV-1 Pol indicating dimeric protein in solution. **c**, SDS-PAGE analysis of fractions from the gel filtration profile of HIV-1 Pol. **d**, Dynamic light scattering analysis of HIV-1 Pol sample shows a generally monodisperse sample.

### Cryo-EM structure determination of HIV-1 Pol

Having optimized the expression and purification of HIV-1 Pol, we used cryogenic electron microscopy (cryo-EM) for high-resolution structure determination. Cryo-EM excels at deciphering structures of dynamic macromolecules and their assemblies, and it is capable of distinguishing different structural states from a single dataset^28^. We collected 901 cryo-EM movies of frozen hydrated HIV-1 Pol using a Thermo Fisher Titan Krios microscope equipped with a K2 direct electron detector (**Extended Data Table 2**). We processed the dataset using an iterative classification procedure, which yielded a final stack of 27,555 particles and a map that was resolved to 3.8 Å within the homogeneous central region composed of globular protein density (**Extended Data Fig. 2-3**). The cryo-EM map enabled the building of an atomic model that was consistent with the experimental density and showed good statistics (**Fig. 2a-b, Extended Data Fig. 3** and **Extended Data Table 2**).

**Figure 2.**
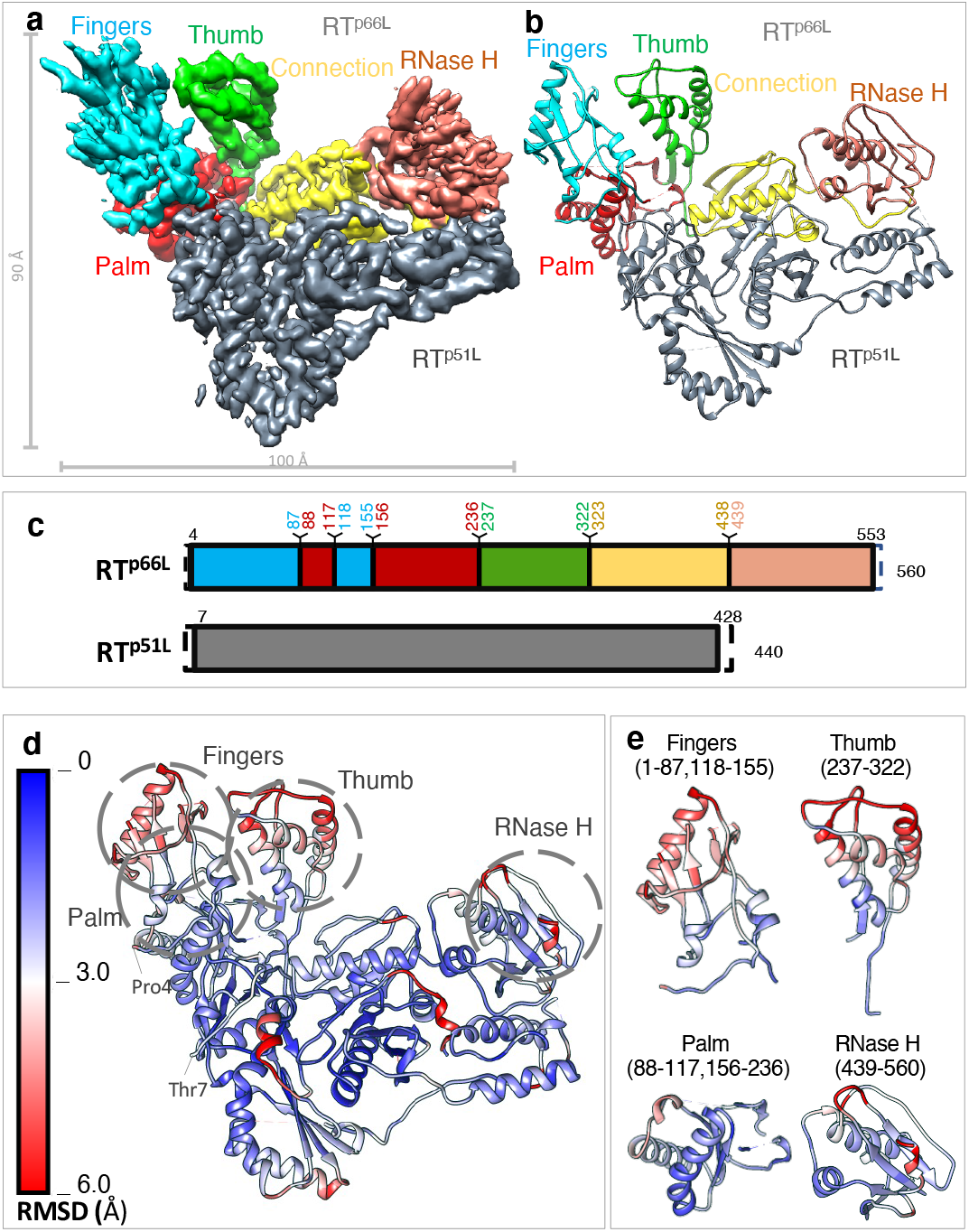
Cryo-EM structure of the HIV-1 Pol dimer. **a**, The high-resolution cryo-EM map of HIV-1 Pol reveals an asymmetric dimer of RT with no additional density for PR and IN at this display threshold. RT^p51L^ is in gray, and RT^p66L^ is colored and labeled by functional domains. **b**, Refined atomic coordinate model labeled and colored accordingly. **c**, Domain architecture for both subunits of RT colored according to **a** and **b**. Full bars represent the first and last residues resolved in the model. Dashed regions represent the regions of the full-length RT that were not resolved in the model. **d**, Root mean square deviation (RMSD) of the high-resolution cryo-EM model compared with mature apo RT (PDB 1DLO). Blue regions represent regions of the model that are structurally conserved between both models, whereas red regions depict portions of the model that are displaced in Angstroms (Å). Functional domains of RT^p66L^ are labeled and indicated by dashed circles. The first and last resolved residues and the buried protease cleavage site (F440/Y441) in the RNase H domain of RT^p66L^ are labeled. **e**, Alignments of the individual subdomains of RT^p66L^ in Pol compared to mature RT p66 show low RMSD within each subdomain.

### Architecture of RT in the Pol Polyprotein

The central core of HIV-1 Pol showed an architecture that is strikingly similar to mature HIV-1 RT (**Fig. 2a-b**). Mature HIV-1 RT is an asymmetric heterodimer containing a p66 subunit, which includes an RNase H domain, and a p51 subunit, from which the RNase H domain has been cleaved by HIV-1 PR between positions 440 and 441. Although the entirety of Pol was present in the sample frozen for cryo-EM analysis, as was confirmed by SDS-PAGE (**Fig. 1c**), only the p66/p51 heterodimer could be clearly resolved within the high-resolution map. This suggested that the remaining polypeptide regions in Pol outside the p66/p51 heterodimer were positionally and/or conformationally variable. The fact that Pol is dimeric in the reconstructed map is consistent with both gel filtration and DLS experiments, both of which suggested that Pol is dimeric in solution (**Fig. 1b, d**). In Pol, the N-terminal residues of the ‘RT p66-like’ (RT^p66L^) and ‘RT p51-like’ (RT^p51L^) subunits are displaced from their usual positions observed in crystal structures of RT (**Fig. 2c**) and extend away from the body of the RT. As will be discussed below, the displacement of these residues can be attributed to the presence of PR upstream of the N-termini of RT^p66L^ and RT^p51L^.

The uncleaved RNase H domain extending from the Pol RT^p51L^ subunit was not visible in our globally refined structures, suggesting that this component has high conformational flexibility. The ordered density for RT^p51L^ ends after residue 428 of the C-terminus of the connection subdomain (**Fig. 2d**). Many of the available crystal structures of HIV-1 RT p66/p51 show weak density in this region and the structure is not usually modeled beyond p51 Q428 (https://www.rcsb.org/uniprot/P03366). The presence of homogeneous density consistent with a heterodimeric p66/p51 structure in Pol suggests that this configuration is produced early in maturation. RT dimerization would stabilize the structure of the RNase H domain in the RT^p66L^, sequestering the F440/Y441 cleavage site in the folded polyprotein and making it inaccessible for cleavage (**Fig. 2d**). Conversely, the F440/Y441 site in RT^p51L^ remains exposed to solvent, as suggested by the lack of density corresponding to this region within the high-resolution cryo-EM reconstruction. The exposure of the F440/Y441 site in RT^p51L^ accordingly likely facilitates the cleavage between F440 and Y441 that creates the C-terminus of p51^29^. Whether the cleavage that frees the IN domain occurs before the RNase H domain of the RT^p51L^ subunit is cleaved is unknown. The structure of the RT heterodimer in Pol means that only one RNase H domain is available for cleavage. This suggests a plausible mechanism for maintaining the stoichiometry of the p66/p51 subunits of RT in virions. Although the density following Q428 in RT^p51L^ does not permit additional residues to be modeled reliably, there is an extension of weak but significant continuous density that threads through a hole in the p66/p51 dimer interface to the side opposite the nucleic-acid binding cleft and into a region that shows unresolved but significant density that could represent the RNase H domain of RT^p51L^. This density provides the first glimpse of the second RNase H domain in RT^p51L^.

### HIV-1 Pol is active for reverse transcription and RNase H cleavage

Given that the p66/p51 heterodimer forms the core scaffold of Pol, we examined whether Pol exhibits either of the enzymatic activities of the mature RT protein: the enzyme (1) copies RNA or DNA using its polymerase domain, and (2) degrades the RNA strand of the RNA/DNA hybrids using its RNase H domain. We found that the pre-formed RT embedded within the Pol polyprotein has both polymerase and RNase H activity on an RNA/DNA template-primer and produces products that are similar to the products produced by mature p66/p51 RT (**Fig. 3a-b**). These data broadly suggest that RT within Pol can carry out reverse transcription. However, when a dsDNA hairpin template-primer aptamer^30^ was used as a polymerase substrate, the pause patterns during DNA synthesis differed between mature HIV-1 RT p66/p51 and the RT embedded in Pol (**Fig. 3c**), suggesting that there are differences in the way the aptamer is engaged by these two RTs. HIV-1 virions containing a PR-RT fusion have been shown to retain PR and RT activities in COS-7 cells, leading to the production of mature infectious virions that were morphologically indistinguishable from WT virions^15,31^. In the RNase H assays we performed, no off-target cleavages were seen, even though two RNase H domains are present in Pol (**Fig. 3b**). These data suggest that the two main enzymatic activities of mature RT are fully recapitulated in the context of Pol.

**Figure 3.**
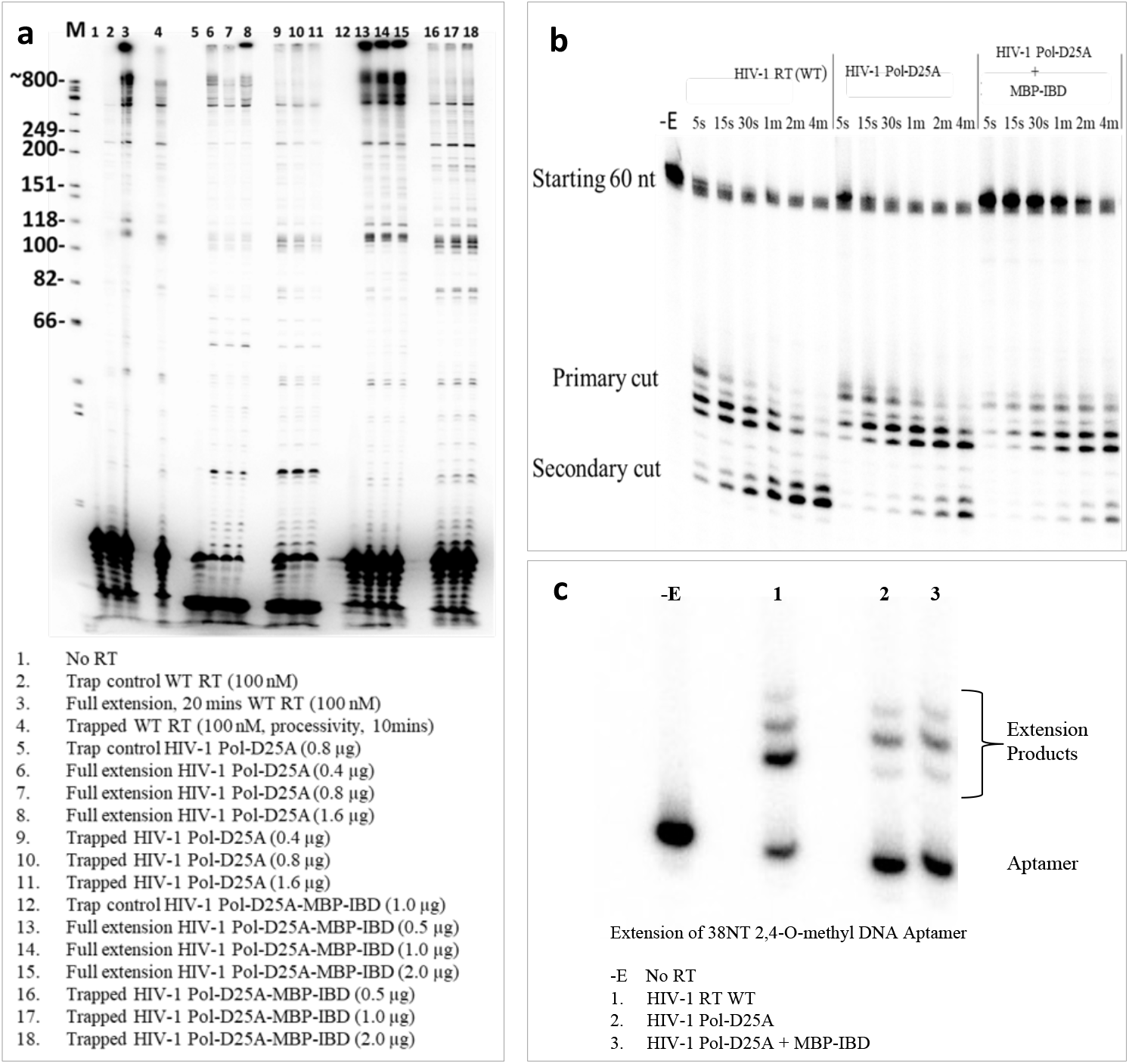
The RT component of HIV-1 Pol is enzymatically active. **a**, Primer extension and processivity assays using an ∼4 kb genomic RNA template (T) derived from the pBKBH10S molecular clone, hybridized to a 5 -^32^P labeled DNA primer (P) (see Methods for details). Heparin was included as a “trap” for the enzymes during the processivity tests. Pol + MBP-IBD indicates the use of a complex between Pol and the integrase-binding domain (IBD) of LEDGF expressed as a maltose-binding protein (MBP) fusion protein. **b**, Analysis of RNase H activity. Positions for the 60-nt starting material and primary and secondary RNA cleavages are indicated. **c**, Extension of DNA aptamer.

### Architecture of PR in the Pol Polyprotein

Class averages derived from the cryo-EM data revealed the presence of indistinct and weak density surrounding the homogeneous region attributed to the core RT p66/p51 heterodimer (**Fig. 4a**). This weak density was present in most of the 2D class averages but was poorly defined in the final reconstructed map (**Fig. 2a** and **Extended Data Fig. 3**). Some of this weak density was located proximal to the N-termini of the fingers subdomains of both RT^p66L^ and RT^p51L^. This density was poorly resolved compared to the RT core in this high-resolution map and was only evident at lower map threshold values. Notably, a global classification approach failed to improve the quality of this weak and likely heterogeneous density. To better resolve this region, we instead employed a focused classification approach that is implemented within the *cis*TEM processing package^32^. This approach works by performing 3-D classification on a region of the map constrained by a spherical mask, while the Eulerian angles and translational shifts remain fixed and aligned with the established orientations used in the reconstruction. The advantage of this approach is that it enables recovery of density for structured regions that are delineated by the boundary of the mask, obviating the need for signal subtraction, a process that can introduce errors if the subtracted density is large (as would be expected if the density assigned to RT was subtracted). The workflow and applications of such approaches have been previously described^33-35^. Using focused classification applied to the region outlined by a spherical mask of radius 35 Å, we found that ∼70% of particles used to generate the high-resolution map contained a mass of density in the expected region for PR, as shown in **Extended Data Fig. 4**. As a control, we also applied a mask with the same radius to a region not expected to contain additional protein mass. As expected, only a negligible amount of density appeared in the mask applied in the control region. The additional density recovered through focused classification within the predominant class (40% of particles, class2) was improved compared to the density that was recovered using global classification alone (**Extended Data Fig. 5**) and was consistent with the size and shape of a PR dimer, which could be unambiguously rigid-body docked into the resulting map (**Fig. 4b-d**). The remaining two classes had either partial or low occupancy and were omitted in the subsequent analysis.

**Figure 4.**
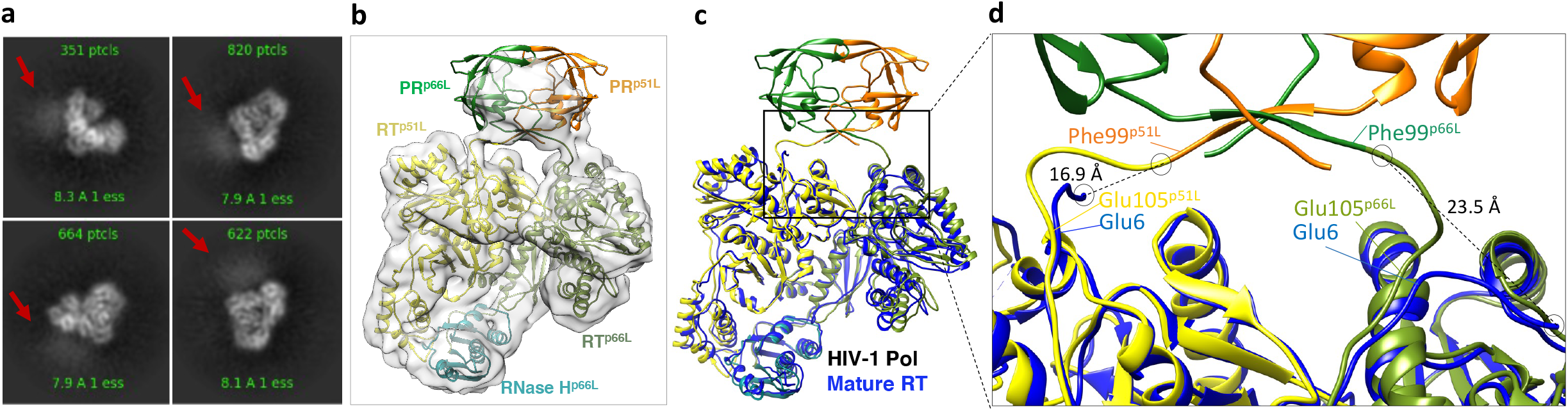
RT dimerization within HIV-1 Pol positions PR into a configuration competent for activation. **a**, 2D class averages of HIV-1 Pol show weak density proximal to RT, depicted by red arrows and suggestive of structural heterogeneity. **b**, A map obtained from focused classification was used for rigid-body docking of mature apo PR dimer (PDB ID 2HB4) and building the residues linking the PR and RT regions of Pol. **c**, Superposition of Pol and apo RT (1DLO, blue). **d**, Close-up view of the PR-RT linkers in Pol. Comparison between the N-termini of RT (1DLO) and the RT portion of Pol shows distinct conformational changes that accompany bringing the PR and RT dimers into proximity. The displacements of the N-terminal residues of RT^p51L^ (∼17 Å) and RT^p66L^ (∼24 Å) compared with the same residues in mature RT (1DLO) are depicted by dashed lines.

The PR component of Pol is directly adjacent to the RT core, connected to the N-terminal residues of the fingers subdomains of both RTs (PR^p66L^ connects to RT^p66L^ and PR^p51L^ to RT^p51L^) of the Pol dimer. The resolution of the PR in our structures (6-8 Å, see **Extended Data Fig. 4**) is lower than the resolution of RT (3.5 Å), which is suggestive of an orientationally dynamic protein. We could unambiguously identify the orientation of the PR dimer within the cryo-EM density, because the linkers that connect the C-terminal residues of PR to RT (4-6 Å resolution for the linkers, see **Extended Data Fig. 4**) are located directly adjacent to the N-termini of the two fingers subdomains within the heterodimeric RT core. These linkers were also clearly apparent within the cryo-EM density after focused classification (**Fig. 4c-d**). The tethering of C-terminal residues of PR to RT pulled the N termini of RT slightly away from their usual positions in crystal structures of the mature enzyme. The N-terminus of RT contains a proline-rich sequence (with six prolines in the first 25 residues) that facilitates the repositioning of the C-termini of PR. The two Pro100s of Pol (which corresponds to Pro1 of mature RT) are pulled 23.5 and 16.9 Å from their positions in the p66 and p51 subunits of mature RT, respectively. Pro103 (which corresponds to Pro4 of mature RT) is displaced ∼6 Å away from the position of equivalent residue in both mature RT subunits, while Pro108 (which corresponds to Pro9 of mature RT) is in approximately the same position in both subunits of Pol and mature RT (**Fig. 4d**). PR is therefore held at “arm’s length”, away from RT, by its own C-terminal residues and the N-terminal residues of RT. There is relatively little buried surface area at the PR-RT junction (253 Å^2^)^36^. This somewhat loose association between PR and RT may contribute to the conformational mobility of the PR relative to RT that gives rise to the weak density we observed (in the focused classification class2). This is likely to be functionally relevant, since the PR and RT must eventually be liberated from Pol during maturation. This configuration exposes the PR/RT cleavage site to solvent (and to PR), making it accessible to cleavage during maturation. Thus, the formation of the highly stable RT dimers appears to be a prelude to the activation of the PR by bringing the monomeric PRs into close proximity, which will, in turn, promote PR dimerization and may relieve some of the destabilizing effects known to be associated with upstream p6* residues, as observed in *in vitro* studies^9^. The heterogeneity observed for PR within the cryo-EM map of Pol may help to explain the sluggishness of the initial cleavage events during maturation as well as the dramatically reduced inhibitory activity of active site PR inhibitors against the immature PR in the context of Gag-Pol^12,37-39^. Taken together, our results reveal that, in the context of Pol, PR is connected by two flexible linkers that loosely tether PR and RT, ensuring that the PR can efficiently dimerize during the maturation stage of the viral replication cycle.

### Arrangement of IN in the Pol Polyprotein

The high-resolution cryo-EM map of Pol at reasonable contour levels only contained density for the RT portion of Pol, despite the protein sample containing the additional PR and IN domains. Focused classification in the region expected to contain PR revealed additional protein density, so we applied this same strategy to find additional density extending from the C-terminal ends of RT within Pol. First, a 21 Å radius mask was applied to the region directly downstream of the RNase H domain of RT^p66L^, into which the IN domain should extend. As a control, a mask with the same radius was applied onto a region that is not expected to be occupied by protein mass. This approach recovered the N-terminal domain (NTD) of IN (residues 1-49) in one class corresponding to ∼20% of the particles from the high-resolution reconstruction (**Extended Data Fig. 6, FC-1**). Importantly, no density appeared after focused classification in the control experiment (**Extended Data Fig. 6, FC-2**). Despite using larger masks and trying different mask placements, we could not determine the remaining density for IN, which should contain the catalytic core domain (CCD) and C-terminal domain (CTD) of IN. This is not surprising, because IN contains two long flexible linkers—one connecting the NTD to the CCD, and the other connecting the CCD to the CTD. As a result, most of the IN density is not visible in our cryo-EM maps, consistent with conformational variability imparted by the presence of flexible domain linkers within IN. The low-resolution cryo-EM density belonging to the IN NTD domain, which was seen with our constructs, emanated from the C-terminus of the RT^p66L^ subunit (**Fig. 2c, Extended Data Fig. 6, FC-1**). RT and IN are connected by a loop that serves as the hinge for IN (**Fig. 1a**). The flexibility of the RT/IN junction, which is solvent-exposed, makes it easily accessible by the PR; cleavage at the junction is required to produce the mature IN protein.

We next performed focused classification downstream of Pol^p51L^ in the hopes of locating the second RNase H domain of the RT^p51L^ chain that will be eventually cleaved to form p51, as well as any additional IN density. Again, a 21 Å radius mask was applied, but this time the mask was placed proximal to the last visible residue of RT^p51L^ and near where the last residue of p51 that is modeled in most crystal structures of mature RT. Focused classification revealed some additional density extending in this region, but it was small and not defined well enough to model any high-resolution structures of the missing protein via rigid-body docking (**Extended Data Fig. 7**). Ultimately, focused classification could only localize the NTD of IN extending from the Pol^p66L^ chain and hint at where the missing RNase H domain of Pol^p51L^ sits within Pol. The remaining domains (the CCD and CTD of IN extending from Pol^p66L^, and the RNase H and IN domains connected to Pol^p51L^) must be either disordered or connected by flexible linkers.

The Pol structure is clearly dimeric, and we only observed a single band on the SDS-PAGE gel after purification (**Fig. 1c** and **Extended Data Fig. 1a**, respectively) which suggested that no cleavage occurred during the preparation of the Pol polyprotein. Thus, both IN protomers must be present in each Pol dimer. Ultimately though, only a single NTD of one of the IN protomers could be resolved by focused classification, and we were unable to establish the multimeric state for IN based on the cryo-EM map. This raised the question of how IN is arranged within Pol. To address this question and investigate the multimeric state of IN, we relied on the fact that the IN-binding domain (IBD) of LEDGF/p75 binds specifically to dimeric and tetrameric HIV-1 IN multimers. We found that, when co-expressed with the IBD of LEDGF/p75 fused to the maltose-binding protein (MBP-IBD), Pol forms a complex with IBD and remains stably bound even after a 1.0 M salt wash on a dextrin sepharose column (**Extended Data Fig. 8a-b**). Because the IBD is known to bind to highly conserved residues at the CCD/CCD dimer interface^40^, we infer that the IN, as part of uncleaved Pol, can form CCD-mediated dimers, at least in the presence of IBD. As dimerization of RT is evident in Pol based on our cryo-EM structure, and focused classification of PR revealed some degree of dimerization for PR, at least some dimerization of IN within Pol seems plausible. Overall, our data suggest that RT and IN dimerization are intricately linked to PR activation.

## Discussion

Collectively, our structural, biophysical, and biochemical data demonstrate that a p66/p51 asymmetric dimer resembling mature RT forms the core scaffold of HIV-1 Pol prior to cleavage by PR and that this complex is enzymatically active in terms of reverse transcription and cleavage of an RNA/DNA duplex. Additionally, our cryo-EM structure indicates that PR can form dimers while part of Pol. Our constructs contain the D25A mutation in the PR, which renders even the mature PR dimer less stable than WT due to disruption of a hydrogen-bonding network that stabilizes the dimer at the fireman’s grip. These observations suggest that RT plays a positive role in holding the PR subunits in close proximity, enhancing their ability to form enzymatically active dimers. Consistent with this, it has been reported that: i) while a minimal TFR-PR D25N construct fails to dimerize, the addition of the first seven amino acids of RT leads to dimerization^38^; ii) inhibition of RT dimerization is deleterious to Gag and Gag-Pol processing during maturation^41^. Conversely, stabilization of RT dimers using molecules such as efavirenz leads to premature PR activation, presumably by enhancing the formation of enzymatically competent PR dimers within the polyprotein^3,42^. We propose a model of PR activation occurring in three general stages (**Fig. 5**). While the first stage has yet to be visualized by solving the Gag-Pol structure (where the Gag portion could play a role in dimerization propensity), the Pol structure presented here may represent an intermediate stage. Hence, the propensity of PR to dimerize—regulated by the Gag:Gag-Pol 20:1 molar ratio and the sequences surrounding PR in the polyprotein—tunes the dimerization and activation of PR and viral maturation.

**Figure 5.**
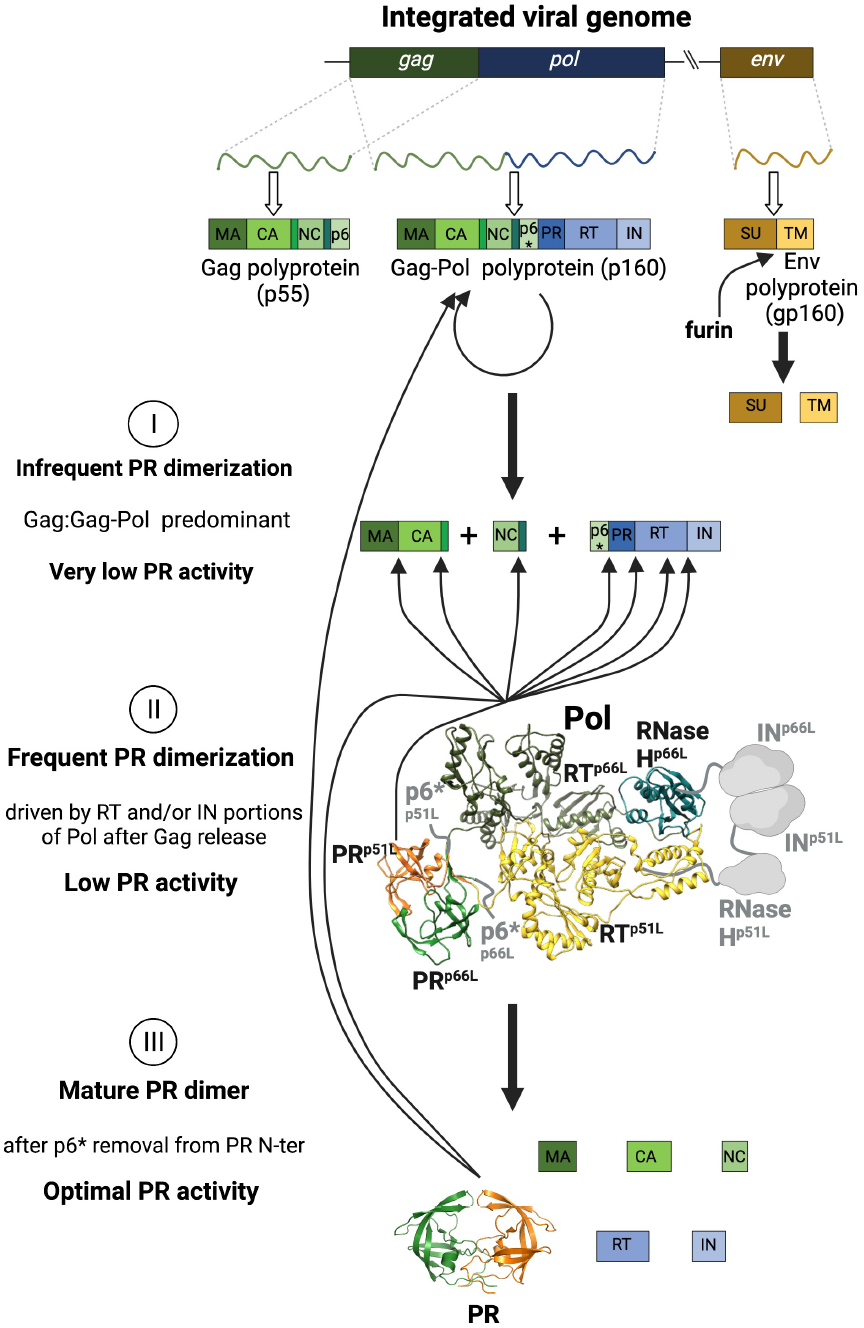
Model of PR activation during HIV maturation. The maturation process is divided into three stages according to the dimerization propensity and activity of PR. **I**. Initially, most Gag-Pol molecules are associated with multimers of Gag, and dimers of Gag-Pol may be infrequent. Additionally, dimers of PR within Gag-Pol may be unstable and necessitate extensive sampling in order to adopt a catalytically active conformation. Accordingly, the PR activity of Gag-Pol is low. **II**. In an intermediate stage, once PR (most likely within a Pol-containing polyprotein precursor) has cleaved Pol from Gag, free Pol predominantly forms dimers through dimerization of the RT portion of Pol as observed in these cryo-EM studies. This increases the dimerization propensity of PR (∼40% particles analyzed by cryo-EM contain PR dimers according to the focused classification results), driven by RT and/or IN domains of Pol. PR activity within Pol may still be suboptimal but is likely higher than within Gag-Pol. **III**. The mature PR dimer should be fully active after p6* removal from the PR N-terminus, increasing processing activity and rapidly completing viral maturation. Created using BioRender (biorender.com).

Of note, the assembly of this dimeric Pol entity could also be affected by the formation of IN dimers. The initial site of nucleation for the formation of the Pol dimer currently remains unknown. This extremely stable association brings two PR monomers, and their respective N-termini, into close proximity, which enables them to form enzymatically active dimers. Thus, RT and IN homodimerization partially offsets the dimerization inhibitory effect of the p6* residues at the N-termini of PR which might not be relieved on its own^38,43^. Consistent with this observation, Hoyte *et al*.^44^ report that not only do IN T124N/T174I mutations weaken the binding of an allosteric inhibitor at the CCD dimer interface, but these mutations also inhibit maturation of HIV-1 virions by impairing Gag and Gag-Pol processing, presumably by inhibiting IN dimerization. In the case of PFV, IN has been shown to be required for the dimerization of Pol and PR activation^45^.

Studies of the proteolytic processing of Gag-Pol in cells suggest that stabilization or destabilization of RT dimerization can lead to either enhanced or diminished PR activity^3^. Our structures imply that RT dimerization will enhance PR dimerization and activation. The two PR monomers are drawn together by the proximity of the N-termini of the two RTs. The integrase (IN) domain is largely not observed in our structures, likely due to flexibility between RT and IN and within the subdomains of IN; however, the ability of Pol to bind the IBD domain of LEDGF/p75 suggests that at least the IBD-binding site in IN is dimeric in Pol. Taken together, our results show how HIV-1 maturation leverages the dimerization interface in the RT portion of Pol to regulate the maturation of polyprotein precursors. This regulation controls the timing of the dimerization and activation of PR, which is slow for isolated PR monomers^9^. The coupling of PR activation to the dimerization of RT helps to delineate a key role for RT in PR activation that was initially suggested by biochemical and virological studies^41,42^. The observation that RT adopts a p66/p51 heterodimer-like conformation in Pol explains why only one RNase H domain is available for cleavage and offers a plausible explanation for the maintenance of the strict 1:1 stoichiometry of p66/p51 in virions.

## Materials and Methods

### Protein expression

HIV-1 Pol constructs of the BH10 strain (gift from Michael Parniak) were cloned into a pET28a vector and transformed into BL21-CodonPlus (DE3)-RIL cells (Agilent Technologies) alone or co-transformed with MBP-IBD. The MBP-IBD construct was cloned into a pCDFDuet vector, compatible with co-transformation with the pET28a HIV-1 Pol vector. In both instances, colonies were selected and inoculated into 100 mL of overnight culture using JJH media (**Extended Data Table 1**), composed of 1.5% tryptone, 1.0% yeast extract, 1.5% NaCl, 1.5% NZ Amine, and 50 mM MgSO_4_ or MgCl_2_ at pH 6.5 in a 500 mL Erlenmeyer flask containing 50 µg/mL kanamycin and 34 µg/mL of chloramphenicol (and 50 µg/mL of streptomycin when co-transforming with MBP-IBD). The culture was shaken at 230-250 rpm overnight at 37 °C, then diluted into 1 L of JJH media with 5% glycerol at pH 6.0 in a 2.5 L Erlenmeyer flask, supplemented with 50 μg/mL of kanamycin (and 50 µg/mL of streptomycin when co-expressing with MBP-IBD). The cells were allowed to grow to an O.D of 2-2.5 at 37 °C and then for at least 1 hour at 15 °C. Subsequently, protein expression was induced with 1 mM IPTG, and culture was grown overnight at 15 °C. 50 mM phosphate buffer at pH 6.0 was added to the media as a buffering agent. Cells were harvested by centrifugation at 4,000xg for 30 minutes, the cell pellet resuspended in 100-150 mL of lysis buffer (100 mM Tris-Cl buffer at pH 8.0, 600 mM NaCl, 0.5% Triton X-100, 10% glycerol, 30 mM imidazole, and 2 mM TCEP) with 1 mM PMSF, 1 µM each of pepstatin A and leupeptin, and cells were sonicated for 10 minutes on ice. The cellular debris were spun down at 38,000xg for 30 minutes.

### Protein purification

The previous supernatant was loaded onto a nickel gravity column pre-equilibrated with the lysis buffer, then washed with: 1) 5-10 column volumes (CV) of lysis buffer, 2) 5-10 CV of high salt buffer wash (1.5 M NaCl) in lysis buffer, 3) chaperone wash of 5 CV (containing 5 mM ATP, 5 mM MgCl_2_, 50 mM imidazole and lysis buffer), and 4) 3 CV wash with the lysis buffer. Protein was eluted with 4 CV of 80 mM Tris pH 8.0, 600 mM NaCl, 500 mM imidazole, 10% glycerol, 2 mM TCEP, and diluted 2-fold with water before loading onto a 5 mL HiTrap heparin column pre-equilibrated with 30 mM Tris-Cl pH 8.0, 300 mM NaCl, 5% glycerol, 1 mM TCEP. The column was washed with loading buffer until the background UV absorption was negligible. Elution of protein was carried out with the loading buffer containing 1 M NaCl, and protein was concentrated and injected onto a Superose 6 Increase 10/30 GL gel filtration column pre-equilibrated with 20 mM Tris-Cl pH 8.0 and 250-300 mM NaCl. Fractions containing pure protein were pooled, concentrated to 0.2-0.5 mg/L, flash frozen and stored at -80 °C. For co-expression with MBP-IBD, glycerol was removed from the nickel column elution buffer, while elution from heparin column was done using 20 mM Tris-Cl, pH 8.0, 1.0 M NaCl, followed by an MBP Trap HP step, washed with the previous buffer and eluted with 20 mM Tris-Cl, pH 8.0, 600 mM NaCl, 10 mM maltose. Gel filtration chromatography suggested a dimeric molecule of Pol in solution at concentrations between 0.2-0.5 mg/mL in both cases (**Fig. 1b-c** and **Extended Data Fig. 8a-b**, respectively), consistent with DLS experiments (**Fig. 1d** and **Extended Data Fig. 8c**, respectively).

### Cryo-EM specimen preparation for electron microscopy

HIV-Pol protein sample at ∼0.4 mg/ml in gel-filtration chromatography buffer (20 mM Tris-Cl pH 8.0 and 250-300 mM NaCl) was applied (2.5 μl) onto freshly plasma treated (7s, 50W, Gatan Solarus plasma cleaner) holey gold UltrAuFoil grid (Quantifoil), adsorbed for ∼1 minute, and plunged into liquid ethane using a manual cryo-plunger operated inside the cold room (∼4 °C).

### Cryo-EM data collection

Data were acquired using the Leginon software^46^, installed on an FEI/Thermo Fisher Scientific Titan Krios electron microscope at the Scripps Research Institute, operating at 300 keV and using a K2 summit direct electron detector operating in counting mode. All data collection statistics are summarized in **Extended Data Table 2**.

### Cryo-EM data processing

Movies were corrected for beam-induced movement using MotionCor2^47^, implemented within the Appion platform^48^. Individual frames were gain corrected, aligned, and summed with the application of an exposure filter^49^. The generated sums excluding the first frame (frames 2-60) were used as input for contrast transfer function (CTF) estimation and particle selection performed in Warp^50^. The particle stack generated in Warp was used as input for 2D and 3D classifications in CryoSPARC^51^, followed by non-uniform (NU) refinement^52^. 2D classification was used to remove particles that did not produce clean class averages. 3D classification using two classes, followed by NU refinement using the particles giving rise to the highest-resolution reconstruction from the two classes, were performed iteratively until no further improvement in resolution and map quality was observed. This reconstruction was then used as input to another round of heterogeneous 3D classification, with the entire stack as input. After another iterative workflow entailing 3D classification and NU refinement, one round of CTF refinement and a final round of NU-refinement was performed (**Extended Data Fig. 2**). The resolution of the RT portion of Pol was assessed using the conventional Fourier Shell Correlation (FSC) analysis^53^, for both masked and unmasked half-map and map-to-model curves, using the “mtriage” tool implemented within the Phenix package^54^. The masks for resolution calculation were automatically generated with mtriage. Directional resolution volumes were generated using the 3D FSC tool^55^, whereas the local resolution was generated using sxlocres.py^56,57^. The final 3D reconstruction was used as input for focused classification performed in cisTEM^32,35^. Different sphere dimensions were tested for focused classification and the final reconstructions used spheres of 35 Å (PR), and 21 Å (RNase H^p51L^, IN, negative controls) radius. The focused classification jobs were performed in manual refinement node without adjustment of either angles or shifts for 30 cycles. For all jobs, we asked for three classes and a global mask of 100 Å was applied.

### Model building and refinement

The model from the RT portion of Pol was built using the initial coordinates from the crystal structure of the apo RT^58^ (PDB 1DLO). The model was aligned to the cryoSPARC sharpened map, and 500 Rosetta_Relax jobs were performed for the relaxation of the backbone bond geometry to improve protein energy landscape for the next step of modeling^59^. Individual regions that diverged from the high-resolution crystal structures were adjusted manually as necessary in Coot^60^. Final models were generated by iterative model building using Coot^61^ and refinement with phenix.real_space_refine^62^ (with secondary structure restraints). The geometry of the final models and other validation statistics were reported by Molprobity^63,64^. The relevant refinement statistics are summarized in **Extended Data Table 2**. High-resolution structural figures were prepared for publication using UCSF Chimera^65^.

### Primer extension and processivity assays

For polymerization assays, reactions were performed using an ∼ 4 kb RNA template made by run-off transcription with T7 RNA polymerase using plasmid pBKBH10S that was cleaved with EcoRI. Plasmid pBKBH10S was obtained through the NIH HIV Reagent Program, Division of AIDS, NIAID, NIH: Human Immunodeficiency Virus 1 (HIV-1) BH10 Non-Infectious Molecular Clone (pBKBH10S DNA), ARP-194, contributed by Dr. John Rossi. After treatment with DNase I to remove the plasmid DNA, isolated RNA was hybridized to a 5 -^32^P labeled DNA primer (5’-GCTTGATTCCCGCCCACCAA-3’) at a ratio of 3:1, primer:template. All reactions were initiated by the addition of dNTPs and MgCl_2_ to other components that were preincubated at 37 °C. For reactions testing processivity, heparin (final concentration 1 µg/µl) was included as a “trap” during the initiation step to sequester enzyme molecules that were free in solution or had dissociated from the primer-template. Trapped reactions were incubated for 10 min and non-trapped for 20 min before terminating with 2X gel loading buffer (90% formamide, 10 mM EDTA pH 8, 0.025% each bromophenol blue and xylene cyanol). Samples were run on a 6% denaturing polyacrylamide gel and visualized with a phosphorimager. For analysis of RNase H activity, reactions were performed in the buffer conditions described above (without dNTPs) using as the substrate for RNase H activity, a 60-nt 5’-^32^P labeled RNA (5’-GGGCGAAUUCGAGCUCGGUACCCGGGGAUCCUCUAGAGUCGACCUGCAGGCAUGCAAGCU-3’) hybridized to a 23-nt DNA (5’-AGGATCCCCGGGTACCGAGCTCG-3’) at 1:2 RNA:DNA ratio as the enzyme substrate (final concentration 10 nM RNA in all assays). Assays were carried out in 80 µl reaction volume at 37 °C and were initiated by the addition of 100 nM (final concentration) p66/51 RT or 1.6 µg of the indicated Pol enzyme. Aliquots were removed at the indicated time points and terminated with 2X gel running buffer, then run on a 10% denaturing gel and visualized with a phosphoimager. For extension of DNA aptamer, reactions were performed in the buffer conditions described in the polymerization assays using a 5’-^32^P labeled 38-nt primer-template mimicking aptamer (5 nM final concentration) that binds HIV-1 RT with pM affinity^30^ as the substrate. The loop-back aptamer has a 5-nt 5’ overhang allowing full extension to 43-nt. Reactions were carried out in 20 µl reaction volume at 37 °C and were initiated by the addition of 10 nM (final concentration) of p66/51 RT, Pol-D25A, or Pol-D25A + MBP-IBD and incubated for 10 min at 37 °C. Terminated reactions were resolved on a 16% denaturing gel and visualized with a phosphoimager.

## Data deposition and availability

The cryo-EM map and atomic model have been deposited into the Electron Microscopy Data Bank and Protein Data Bank under the following accession codes: HIV-1 Pol globally refined with resolved RT heterodimer (EMD-25074 and PDB-7SEP).

## Acknowledgements

The authors thank the following individuals for helpful discussions: Stephen Hughes, Steve Tuske, Sanjay Dey, Ruchi Yadav, Youngmin Jeon, Michael Parniak, Jason Kaelber, Alan Engelman, Mamuka Kvaratskhelia, Joseph Marcotrigiano, Bruce Torbett, Celia Schiffer, and Ron Swanstrom. We are grateful for funding from the following sources: NIH grants U54 AI150472 awarded to DL and EA, R01 AI027690 awarded to EA, and R01 AI136680 awarded to DL. The molecular graphics and analyses were performed with the USCF Chimera package (supported by NIH P41 GM103311). J. J. E. K. Harrison is thankful to the International Institute of Education (IIE) and the Fulbright program for a graduate fellowship award. DL is also grateful to the Margaret T. Morris Foundation.

## Author contributions

J.J.E.K.H., D.L., and E.A. conceived the work and designed the experimental strategies. J.J.E.K.H. developed the media and carried out the production and purification of the protein samples. J.D.B. and L.T. carried out Western blot experiments, media composition analysis and discussions of experimental procedure. J.J.D. performed the activity assays. D.O.P. and J.F.B. carried out cryo-EM data acquisition, image processing, and structure determination. D.O.P. built and refined the models. J.J.E.K., D.O.P., J.F.B., F.X.R., D.L., and E.A. carried out data analysis. J.J.E.K.H., D.L., and E.A. wrote the original draft of the manuscript, and all authors contributed to writing and editing. E.A. and D.L. provided funding and supervised the work.

## Competing interests

The authors declare no competing interests.

Correspondence and requests for materials should be addressed to Eddy Arnold and Dmitry Lyumkis.

## Extended Data Figure Legends

**Extended Data Figure 1.**
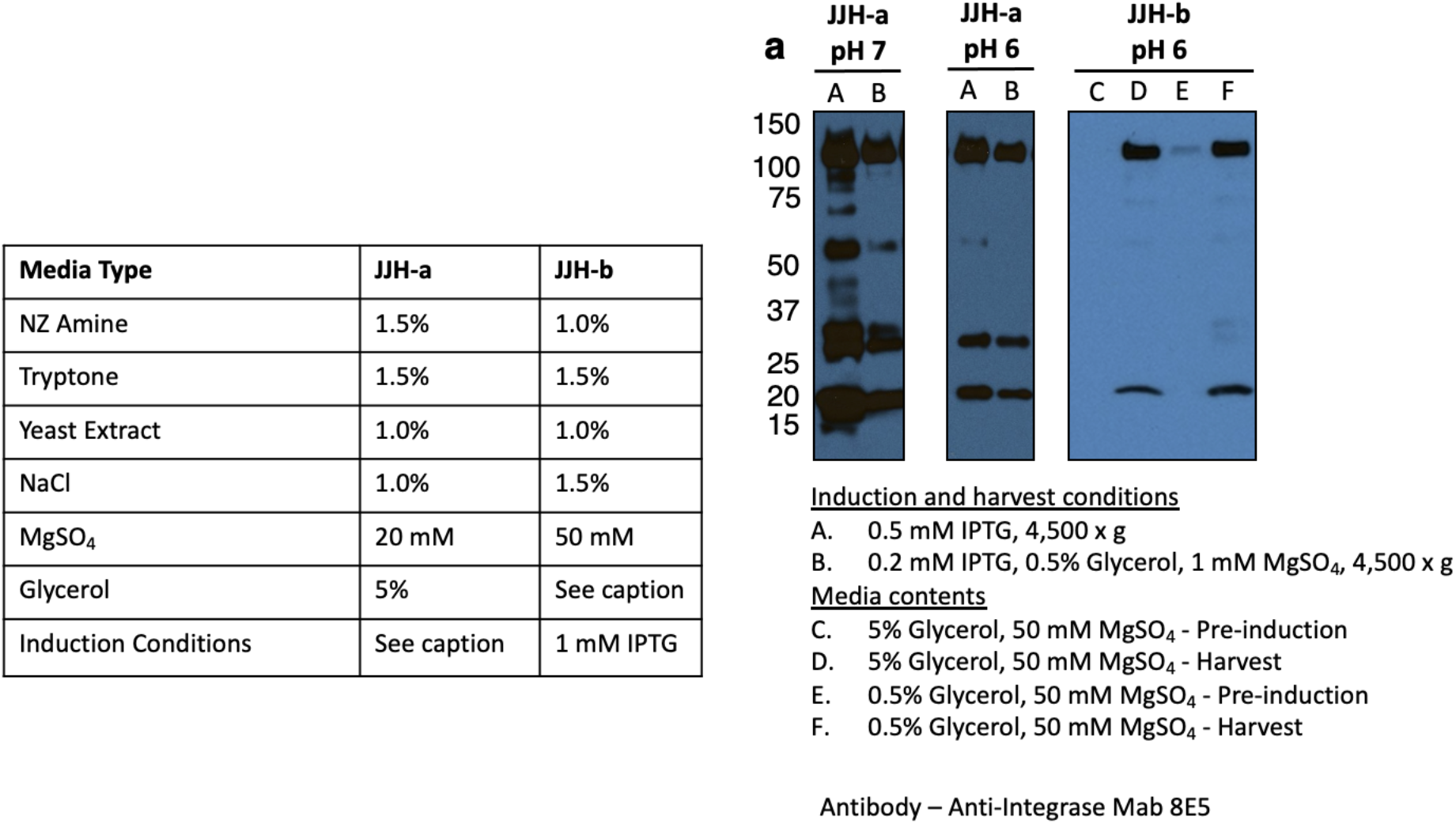
Bacterial expression optimization. Proteolytic degradation of HIV-1 Pol expressed in *E. coli* under different conditions was assessed by Western blot analysis probed with an anti-IN primary antibody 8E5. Intact HIV-1 Pol in our construct has a molecular mass of 119 kDa and any lower molecular weight bands are likely degradation products. JJH is a modified form of LB media (see **Extended Data Table 1**) and has been modified as indicated in this figure.

**Extended Data Fig. 2.**
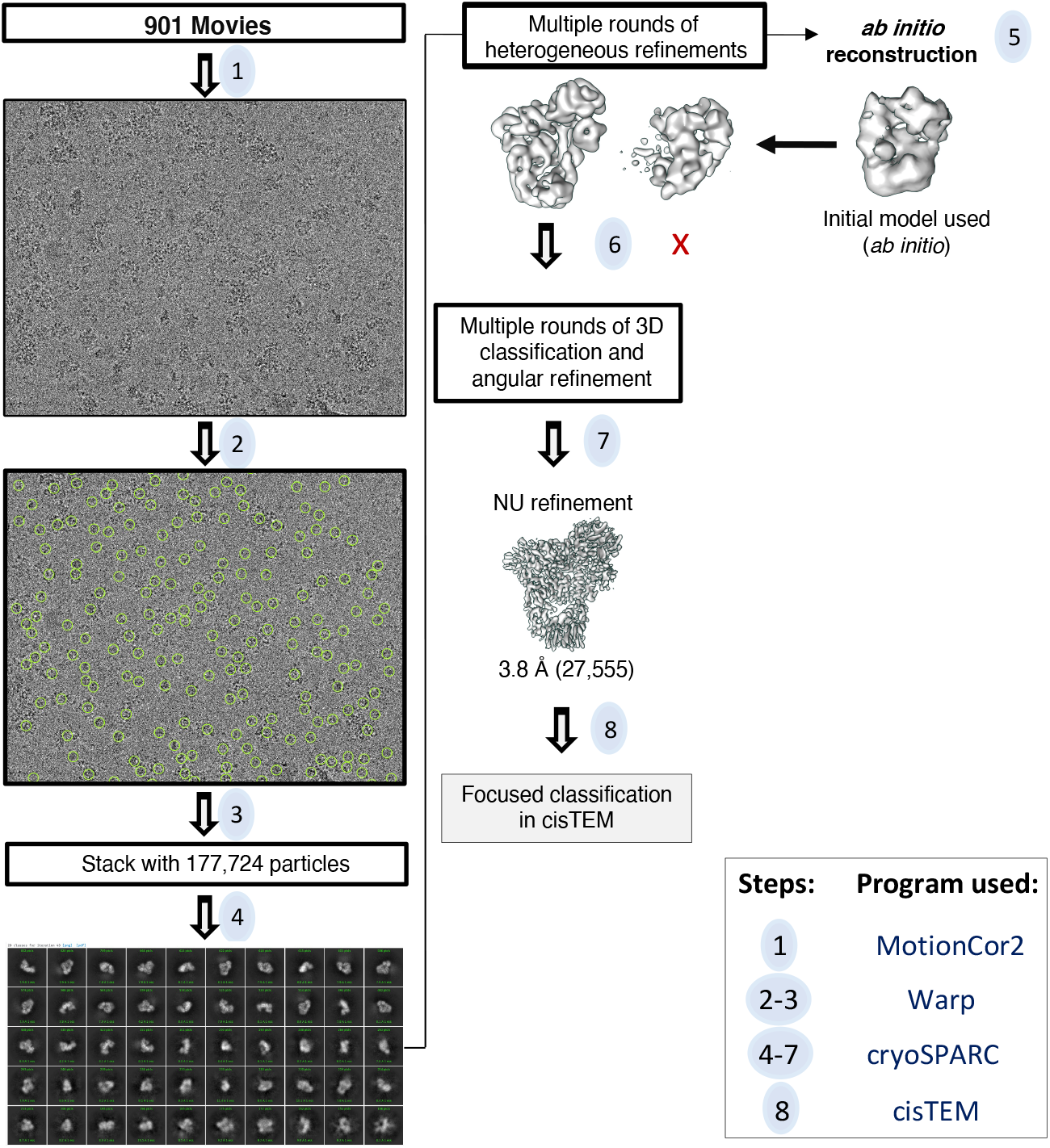
Cryo-EM workflow for HIV-1 Pol. Movies were aligned using MotionCor2 (1). Dose-weighted sums were transferred to Warp for CTF estimation and particle selection (2-3). The stack of particles from Warp were imported into cryoSPARC for 2D classification, *ab initio* reconstruction, and consecutive rounds of 3D classification and refinement (4-5). The final 3D reconstruction from this step was used to retrieve more good particles from the stack and improve the quality of the reconstruction (6). Finally, Global CTF refinement followed by NU-Refinement in cryoSPARC further improved the resolution (7). The final reconstruction from this process was then used as the input for focused classification analysis with the mask in various locations (8; see **Extended Figures 4, 6 and 7**).

**Extended Data Fig. 3.**
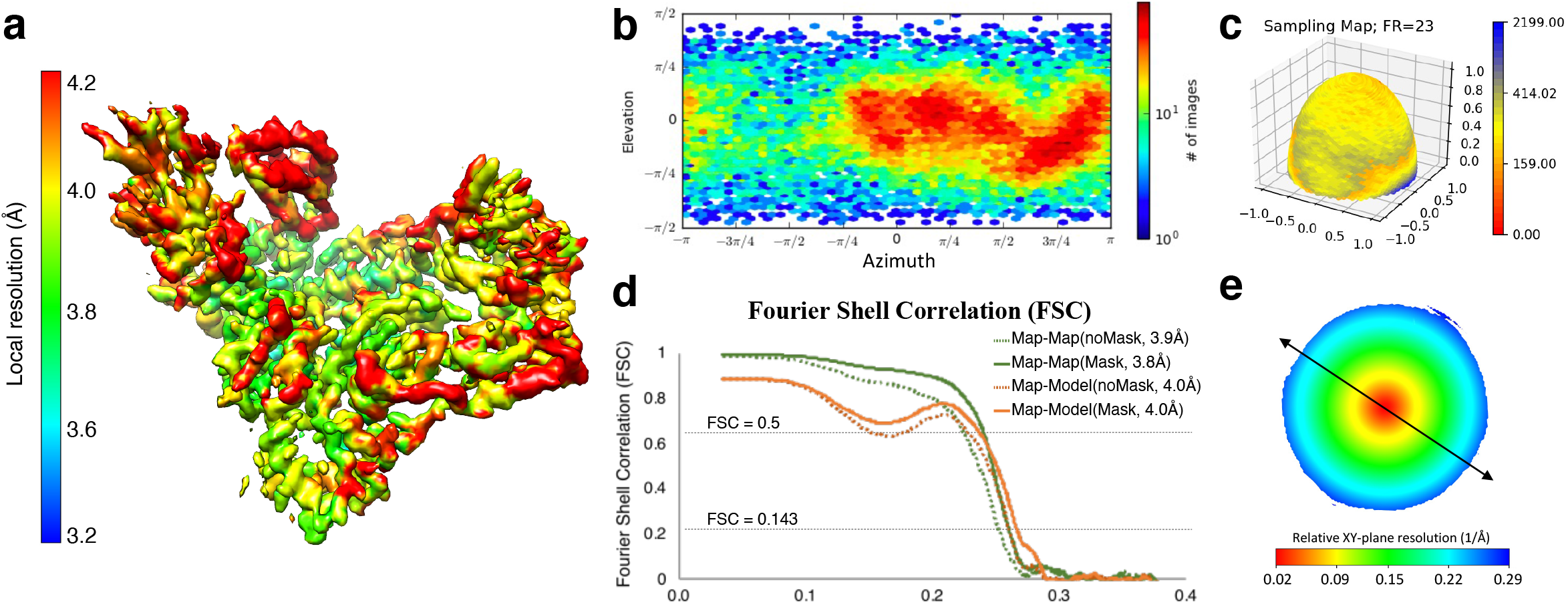
Validation of the cryo-EM map and atomic model of HIV-1 Pol. **a**, The RT portion of Pol, colored by local resolution. **b**, Euler angle distribution plot showing particle orientations contributing to the reconstruction. **c**, Sampling compensation factor (SCF)^66^ depicting the overall angular distribution of the particles comprising the final reconstruction. **d**, Fourier Shell correlation (FSC) for half-map unmasked and masked (green lines dashed and full, respectively) and map-model unmasked and masked (orange lines dashed and full, respectively) analysis. **e**, 3DFSC isosurface plot^55^ showing the resolution range of the map for the least favorable cross-sectional view (values refer to resolution). Arrow indicates the direction of preferred orientation (Z-resolution with respect to the electron beam).

**Extended Data Fig. 4.**
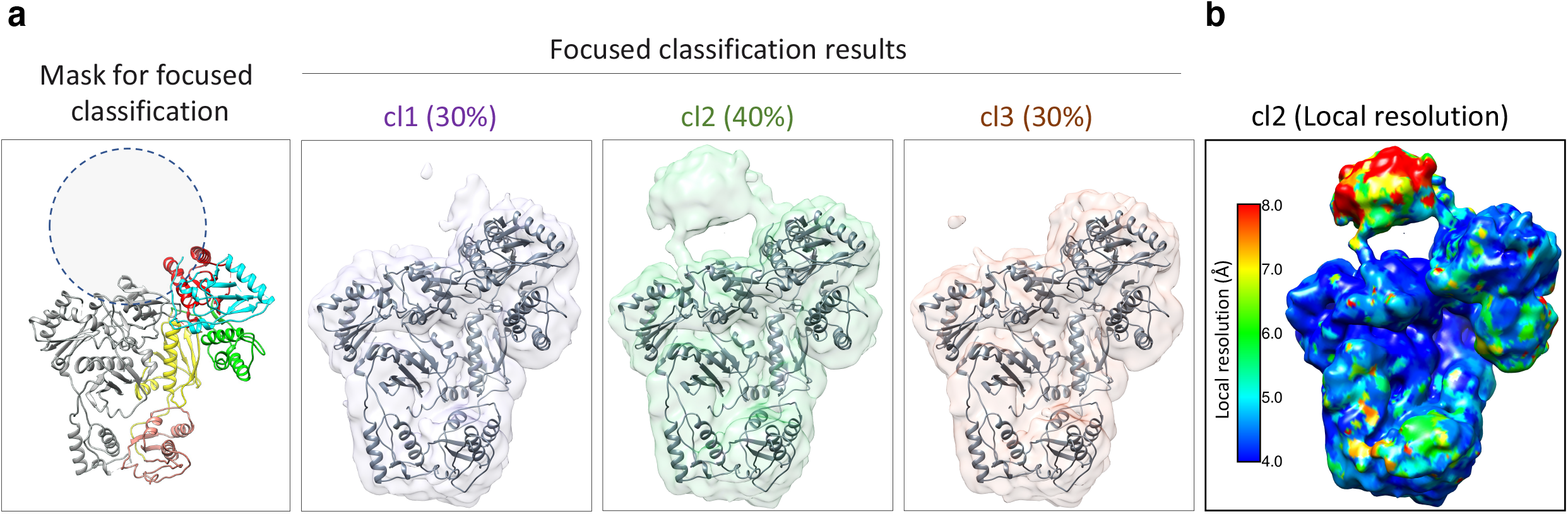
Focused classification of the PR domain within Pol. **a**, A 35 Å radius mask was applied to the region indicated by a dashed circle. This region is proximal to the N-terminal residues of the RT portion of Pol in our high-resolution cryo-EM map. Selection of this area for focused classification was further supported by the presence of weak density in this area within 2D class averages of Pol (**Fig. 4a**). Focused classification was performed in *cis*TEM^32^. Particles were sorted into three classes and the percentage of particles sorted into each class is indicated. Class 2 (F1-2, in green) recovered recognizable density for PR and the linkers connecting PR to RT were completely resolved. This class average corresponds to 40% of the particle making up the high-resolution cryo-EM reconstruction. **b**, the class 2 reconstruction is filtered and colored by local resolution.

**Extended Data Fig. 5.**
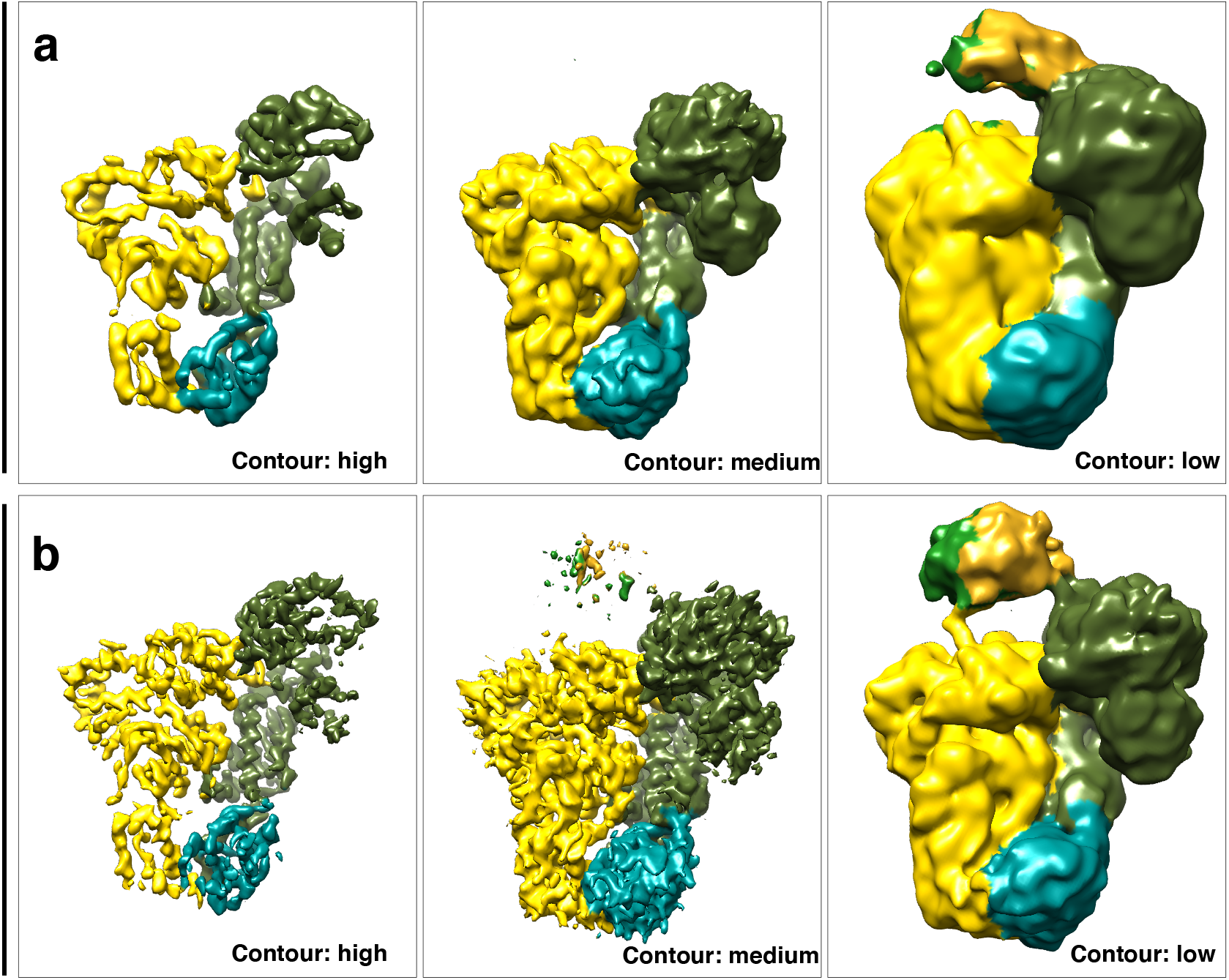
Visualization of PR density at different contour filtering levels. Selected reconstruction generated in *cis*TEM by either **a**, global classification or **b**, focused classification (cl2), depicted at different contour levels in Chimera. The maps are colored according to subdomains.

**Extended Data Fig. 6.**
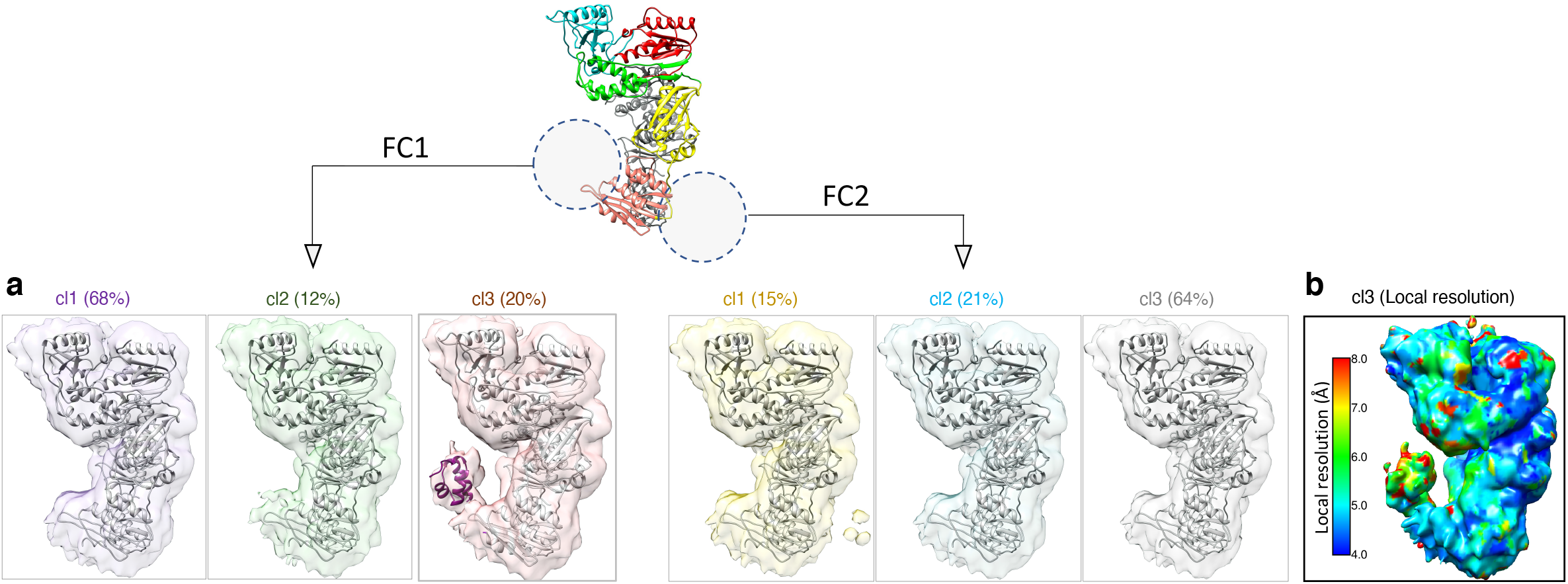
Focused classification on IN density extending from the RT^p66L^ portion of Pol. A 21 Å radius mask was applied to two regions represented by dashed circles: the first was proximal to the C-terminus of RT^p66L^ (FC1) and the second was a control mask placed on the opposite side of the RNase H domain (pink) of RT^p66L^ (FC2). Class 3 containing 20% of the particles in FC1 recovered density that could accommodate the IN N-terminal domain (NTD). The structure of the NTD of IN was docked as a rigid body in this density (violet, PDB ID 6PUT). The negative control classification either did not yield appreciable density or yielded weak discontinuous density when displayed at the same threshold. The application of masks with larger diameters did not yield additional density for the remaining domains of IN.

**Extended Data Fig. 7.**
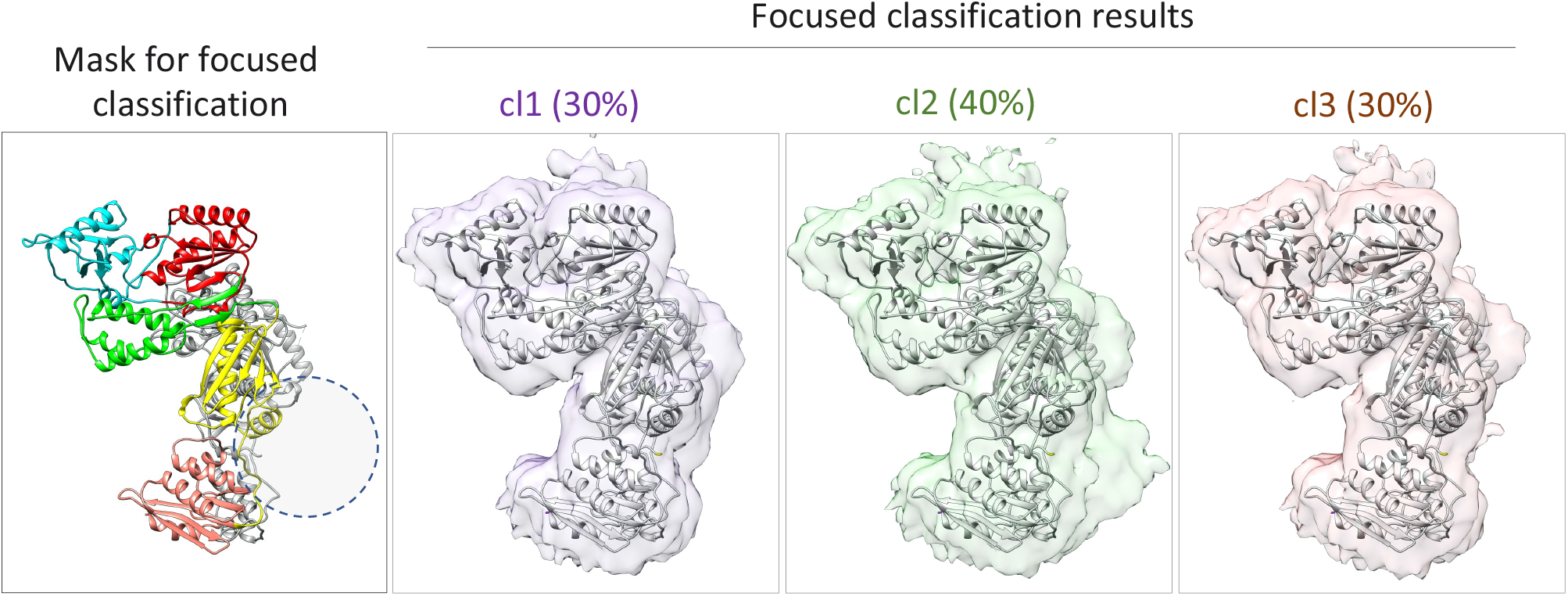
Focused classification of the RNase H domain extending from the RT^p51L^ portion of Pol. A 21 Å radius mask was applied to the high-resolution Pol map proximal to the last visible residues of RT^p51L^ (panel on the right, mask location is represented by a dashed circle). Class 2 corresponds to 40% of the particles and exhibits some additional, though poorly defined density (green). This suggests that there is additional protein density in this region, but that it is mobile and/or partially unstructured.

**Extended Data Fig. 8.**
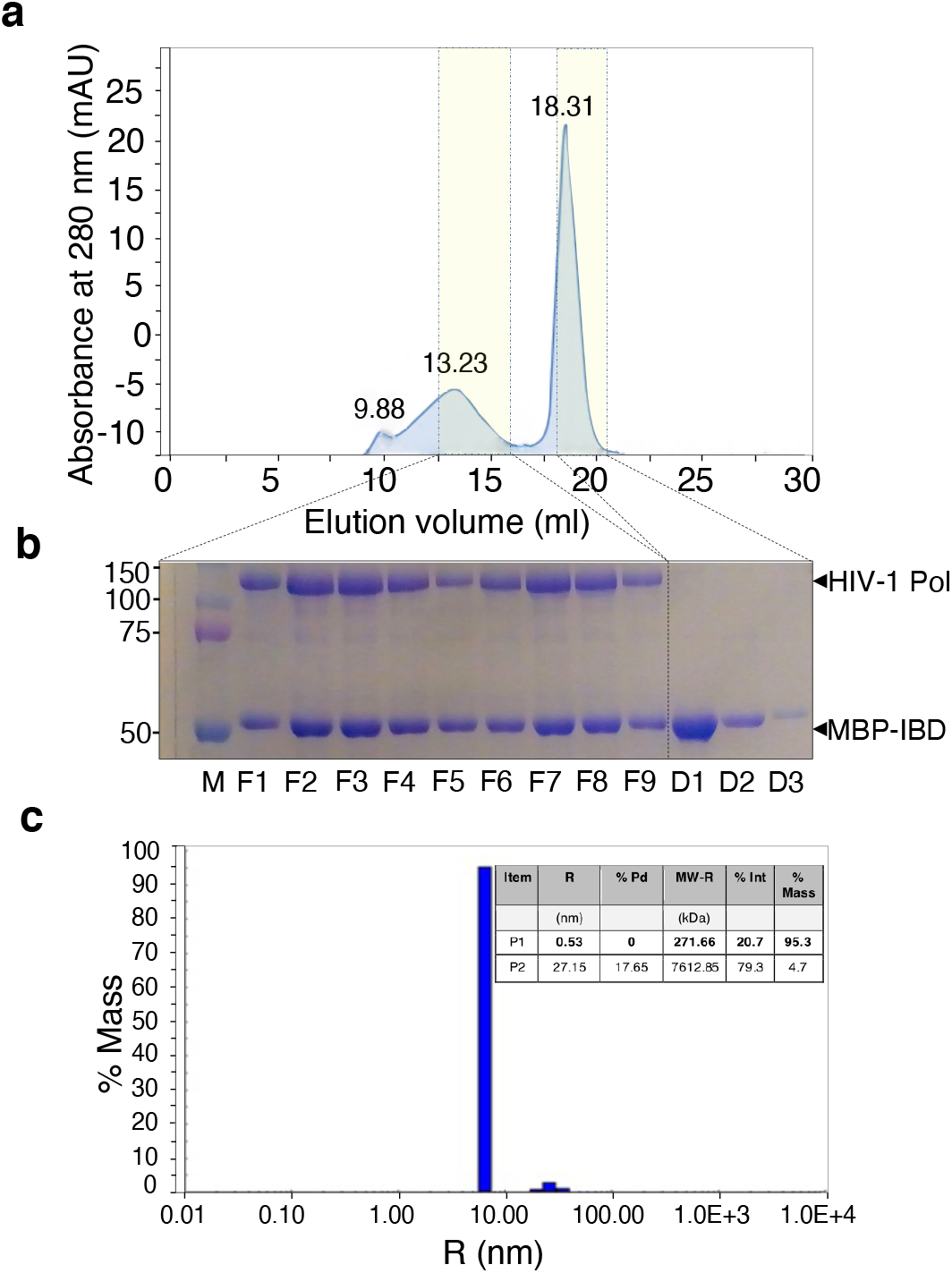
Biochemical and biophysical characterization HIV-1 Pol + MBP-IBD. **a**, Superose 6 increase gel filtration profile of purified HIV-1 Pol + MBP-IBD. **b**, SDS-PAGE gel of fractions from the gel filtration experiment showing good purity of HIV-1 Pol and MBP-IBD. **c**, DLS profile of HIV-1 Pol + MBP-IBD complex showing that the majority species (95% by mass) has a hydrodynamic radius of 6.5 nm and low polydispersity (0%).

**Extended Data Table 1.** Comparison of standard Luria Broth (LB) with Modified LB Media (JJH Media) used to express HIV-1 Pol in *E. coli*.

|  | **LB Media** | **Modified LB Media (JJH Media)** |
| --- | --- | --- |
| Tryptone | 1.0% w/v | 1.5% w/v |
| Yeast extract | 0.5% w/v | 1.0% w/v |
| NZ Amine | - | 1.5% w/v |
| NaCl | 1.0% w/v | 1.5% w/v |
| Glycerol | - | 5.0% w/v |
| MgCl<sub>2</sub> or MgSO<sub>4</sub> | - | 50 mM |
| pH | 7-7.4 | 5.5-6.5 |

**Extended Data Table 2.** Cryo-EM data collection and modeling statistics.

| **Data collection and processing** | **HIV-Pol** |
| --- | --- |
| Microscope | Titan Krios |
| Detector | Gatan K2 |
| Recording mode | Counting |
| Voltage (kV) | 300 |
| Magnification | 29,000 |
| Total dose (e-/Å<sup>2</sup>) | 56.68 |
| Dose rate (e-/Å<sup>2</sup>/s) | 4.72 |
| Defocus range (μm) | -1.0 to - 2.5 |
| Pixel size (Å) | 1.015 |
| Number of frames/movie | 60 |
| Exposure time (sec) | 12 |
| Frame rate (frames/sec) | 5 |
| Symmetry imposed | C1 |
| Number of micrographs used (no.) | 901 |
| Total picked particles (no.) | 177,724 |
| Particles used for final map (no.) | 27,555 |
| Map resolution (FSC 0.143, Å) | 3.8 |
| Local resolution range (sxlocres, Å) | 3.2 - 4.2 |

| **Model statistics** | **HIV-Pol** |
| --- | --- |
| Refinement package (real space refinement) | Phenix |
| Model composition |  |
| Protein (residues) | 930 |
| DNA (nucleotides) | 0 |
| Ligands | n/a |
| Map CC | 0.78 |
| FSC map-to-model (0.5) | 4 |
| RMSD [bonds] (Å) | 0.003 |
| RMSD [angles] (°) | 0.631 |
| All-atom clashscore | 9.7 |
| Molprobity score | 1.81 |
| Ramachandran plot |  |
| Favored (%) | 95.72 |
| Allowed (%) | 4.28 |
| Outliers | 0 |
| Rotamer outliers (%) | 0 |
| CaBLAM outliers (%) | 1.68 |
| C-β deviations (%) | 0 |
| EM-Ringer Score | 2.26 |
| Average B-factor, protein ($\text{\AA}^2$) | 73.92 |
| Average B-factor, DNA ($\text{\AA}^2$) | n/a |

